# Cyanate – a low abundant but actively cycled nitrogen compound in soil

**DOI:** 10.1101/2020.07.12.199737

**Authors:** M. Mooshammer, W. Wanek, S. H. Jones, A. Richter, M. Wagner

## Abstract

Cyanate (NCO^-^) can serve as a nitrogen and/or carbon source for different microorganisms and even additionally as an energy source for autotrophic ammonia oxidizers. Despite the widely distributed genetic potential for direct cyanate utilization among bacteria, archaea and fungi, the availability and environmental significance of cyanate is largely unknown, especially in terrestrial ecosystems. We found relatively low concentrations of soil cyanate, but its turnover was rapid. Contrary to our expectations, cyanate consumption was clearly dominated by biotic processes, and, notably, cyanate was produced *in-situ* at rates similar to that of cyanate formation from urea fertilizer, which is believed to be one of the major sources of cyanate in the environment. Our study provides evidence that cyanate is actively turned over in soils and represents a small but continuous nitrogen/energy source for soil microbes, potentially contributing to a selective advantage of microorganisms capable of direct cyanate utilization.

**One-sentence summary:** Cyanate represents a small but continuously available nitrogen source for soil microbes, contributing to a selective advantage of microorganisms capable of direct cyanate utilization.

## Introduction

Cyanate (NCO^-^) is an organic nitrogen compound that has mainly been of interest in medical science due to its negative effect on protein conformation and enzyme activity (e.g., *1*), in chemical industry as industrial feedstock, and in industrial wastewater treatment, where it is produced in large amounts, especially during cyanide removal (e.g., *2, 3*). However, in recent years, cyanate received more attention in marine biogeochemistry and microbial ecology, with the discovery of the involvement of cyanate in central nitrogen (N) cycling processes, namely in nitrification and anaerobic ammonia oxidation (anammox) (*4, 5*). Despite the emergent recognition of the role of cyanate in marine ecosystems (*6–11*), the environmental role and significance of cyanate in terrestrial ecosystems remain entirely unknown.

It has been shown that cyanate can serve as the sole N source for microorganisms that encode the enzyme cyanase (also known as cyanate hydrolase or cyanate lyase; EC 4.2.1.104) (*8, 12, 13*). This enzyme catalyzes the decomposition of cyanate in a bicarbonate-dependent reaction yielding carbamate, which spontaneously decarboxylates to ammonia and carbon dioxide (*14*). The resulting ammonia and carbon dioxide can then be assimilated (*13*). The enzyme was first discovered in *Escherichia coli* (*15*) and genes encoding homologous proteins have been found since in genomes of various bacteria, such as proteobacteria and cyanobacteria, as well as in archaea, fungi, plants and animals (*16–18*). Cyanase, and thus the potential to use cyanate as a N source, therefore seems to be widespread among prokaryotes and eukaryotes. Generally it is assumed that the main role of cyanase is cytoplasmic detoxification. Cyanate is harmful because isocyanic acid (HCNO), the active form of cyanate, reacts with amino and carboxyl groups, and consequently carbamoylates amino acids, proteins and other molecules, thereby altering their structure, charge and function (*19*). Furthermore, a regulatory function of cyanase in arginine biosynthesis has been proposed (*17*).

Recently, a new physiological role for the enzyme cyanase was described in the chemoautotrophic ammonia-oxidiser *Ca.* Nitrososphaera gargensis. This archaeon encodes a cyanase and was shown to effectively use cyanate not only as a source of N for assimilation but also as a source of energy and reductant (*4*). Moreover, the marine anammox *Ca*. Scalindua profunda as well as several *Ca.* Scalindua single amplified genomes from the Eastern Tropical North Pacific anoxic marine zone also possess a cyanase and it has been suggested that cyanate thus can be directly used as a substrate by anammox organisms (*5*). Cyanate can be either directly utilized by cyanase-positive microorganisms or indirectly by other microorganisms that may assimilate ammonia released by the former. A special case of indirect use of cyanate was shown recently among nitrifiers exhibiting a reciprocal feeding relationship that enables growth of both partners on cyanate. Cyanase-positive nitrite-oxidizers convert cyanate to ammonia, providing the substrate for cyanase-deficient ammonia oxidizers that oxidize ammonia to nitrite, providing, in turn, the substrate for nitrite-oxidizers (*4*).

Cyanate can be formed by photooxidation or chemical oxidation of hydrogen cyanide (*20*), or by hydrolysis of thiocyanate (*21*). Recently, it has also been shown that cyanate is formed in diatom cultures, indicating a biological source of cyanate (*22*). Within living organisms, cyanate may result from the non-enzymatic decomposition of carbamoyl phosphate, a precursor for nucleotide and arginine biosynthesis (*23, 24*). Moreover, urea spontaneously dissociates in aqueous solution, forming cyanate and ammonium (*25*). As urea is the most widely used agricultural N fertilizer worldwide (*26*), it is possibly one of the most significant sources of cyanate in soils on a global scale.

Despite the potential relevance of cyanate as a N and energy source for microorganisms, environmental cyanate sources, concentrations and fluxes (i.e., the production and consumption) are largely unknown, especially in terrestrial ecosystems. Here, we investigated, for the first time, cyanate availability and dynamics in terrestrial ecosystems. We analyzed soil cyanate concentrations across different soil and land management types along a continental gradient and discuss the abiotic behavior of cyanate in the soil environment that controls its availability. We developed a method for compound-specific isotope analysis of cyanate that allowed us to assess biotic and abiotic cyanate turnover processes. To yield further insights into the production and consumption of cyanate in soils, we assessed quantitively the contribution of urea to soil cyanate formation, by combining empirical and modelling approaches that yielded estimates of gross rates of cyanate transformations in soils.

## Results and Discussion

### Cyanate concentrations and the influence of soil pH on its recovery and availability

As cyanate concentrations have not yet been determined in soils, we tested three commonly used soil extractants: water (Ultrapure Water, resistivity >18.2 MOhm), 10 mM CaSO_4_, and 1 M KCl (Fig. 1). If cyanate is strongly adsorbed in soils, increasing salt concentrations of the extractant result in a higher recovery of cyanate. For an alkaline grassland soil (soil pH = 8.3), we found that the recovery of added cyanate was complete for all extractants (i.e., no significant difference between added and recovered cyanate, *t*-test, P > 0.05). However, the recovery of added cyanate differed between extractants for a forest soil with a soil pH of 7.0 (one-way ANOVA, F2,9 = 308.5, P < 0.001). When using 1 M KCl for this soil, recovery was complete (101.5% ± 1.3 SE), whereas the use of 10 mM CaSO_4_ or water resulted in significantly lower recoveries of 85.8% (± 0.7 SE) and 59.5% (± 1.5 SE), respectively. In contrast to the alkaline and neutral soil, cyanate recovery in an acidic grassland soil was on average only 7% for all extractants. For the following experiments we chose 1 M KCl as the extractant, as its extraction efficiency was the same or higher as the others.

**Fig. 1.**
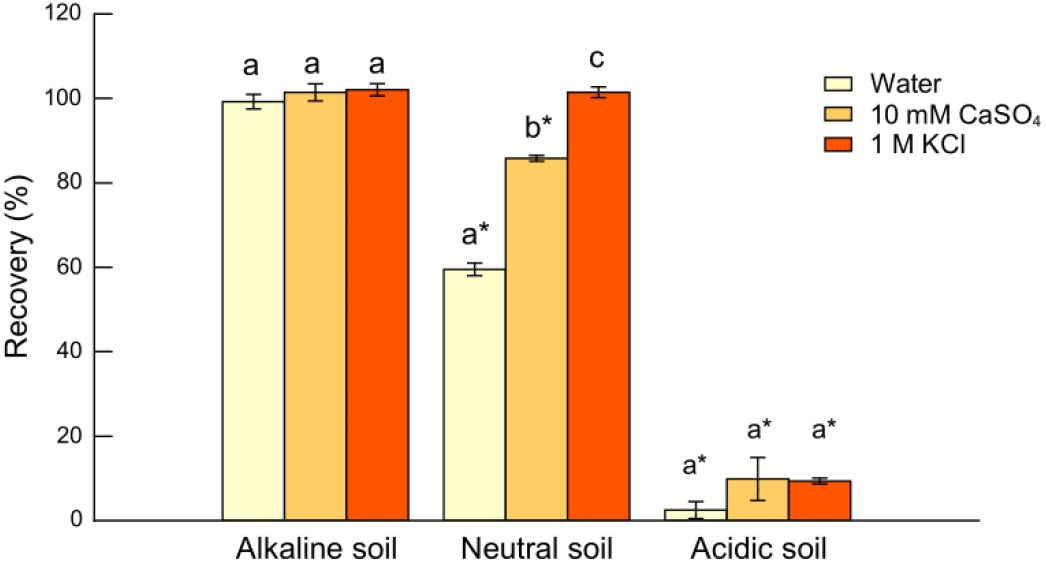
Comparison of extractants for determination of soil cyanate concentration. Cyanate recovery was assessed by spiking the extraction solution with potassium cyanate (final concentration of 15 nM). Three extractants (water, 10 mM CaSO4 and 1 M KCl) were tested for three soils: an alkaline grassland soil (soil pH = 8.3), a pH-neutral mixed forest soil (soil pH = 7.0) and an acidic grassland soil (soil pH = 4.3). Letters denote significant differences between extractants within each soil type (one-way ANOVA followed by Tukey’s HSD test). Asterisks indicate significant differences between added and recovered cyanate (t-test).

To obtain representative data on soil cyanate concentration, we analyzed 46 soils across different soil and land management types along a European climatic gradient (Fig. 2a). Although we used the most sensitive analytical method available to date, with a detection limit in the low nanomolar range in solution (*27*), cyanate was detectable only in 37% of the soils tested (Fig. 2b). Average concentration of soil cyanate was 33.6 (± 8.1 SE) pmol g^-1^ soil d.w., excluding samples below detection limit. Notably, we found that above soil pH 5.7 in 0.01 M CaCl_2_ or pH 6.6 in water cyanate was detectable in all samples, indicating that soils with high pH have higher cyanate concentrations, as also shown by the extraction test mentioned above.

**Fig. 2.**
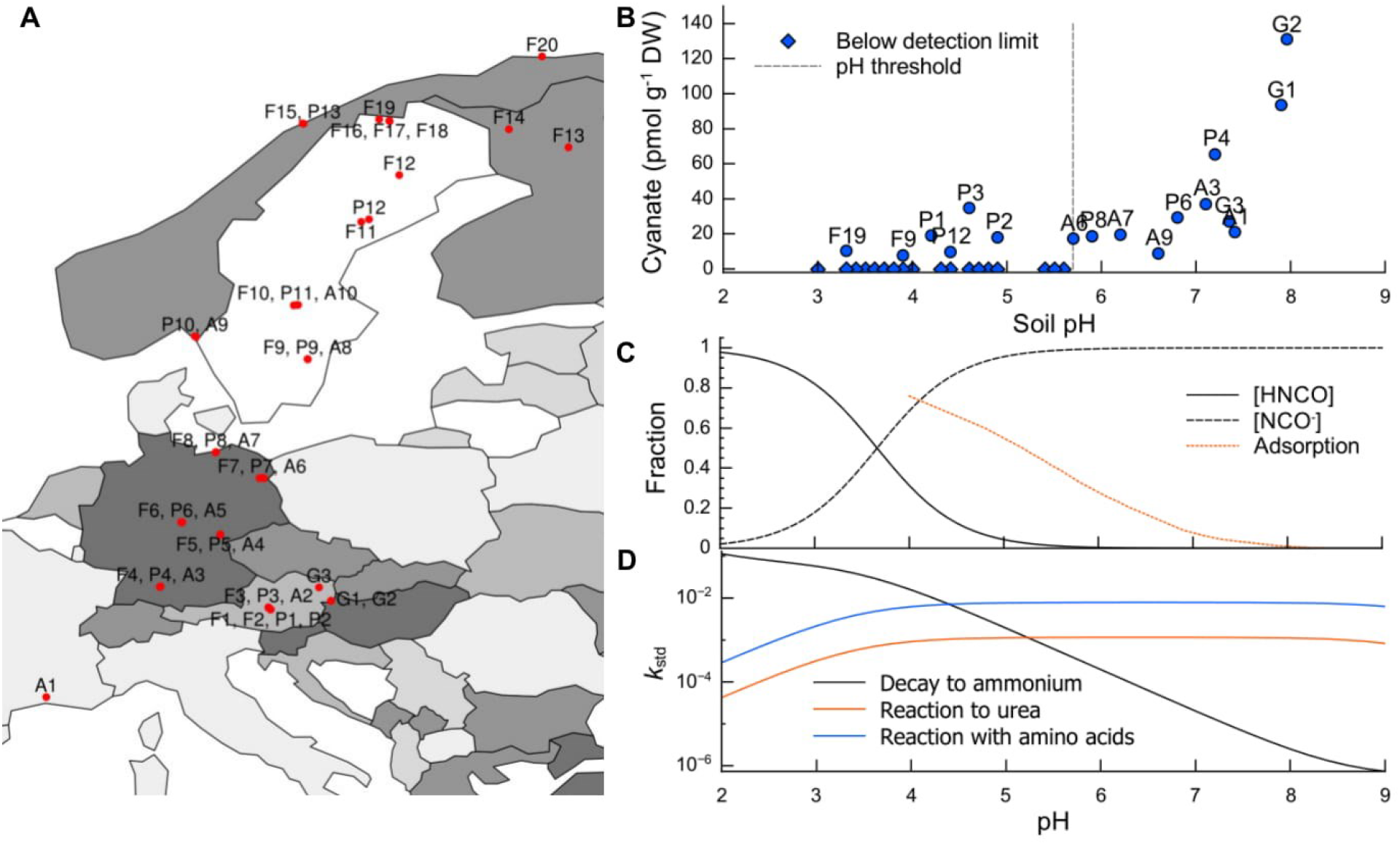
Soil cyanate concentrations and abiotic reactions of cyanate. (**A**) Map of Europe displaying the 46 soil sampling sites: G, grassland; F, forest; P, pasture; A, arable. (**B**) Soil cyanate concentrations (extracted using 1 M KCl) plotted as a function of soil pH in 0.01 M CaCl_2_. The dashed line denotes the soil pH threshold above which cyanate was detectable in all soil samples. (**C**) Acid-base dependency of cyanate and isocyanic acid as a function of pH (HNCO ⇄ H^+^ + NCO^-^; p*K*_a_ = 3.66 at 20°C). The orange dotted line shows the predicted adsorption isotherm of a 10^-4^ M cyanate solution on hydrous ferric oxide (a major component of soil influencing stabilization of compounds) as a function of pH (redrawn from *49*) The equilibrium surface complexation constant was estimated based on correlations of acidity constants and surface complexation constants fitted to adsorption data for other inorganic ions (*28*). (**D**) Standardized rates (*k*_std_; at 20°C) of combined abiotic cyanate/isocyanic acid decomposition to ammonium (equations 6–8, rate constants from equations 9–11), the reaction of cyanate with ammonium to urea (equation 3, rate constants from equation 5) and the reaction of isocyanic acid with the amino group of glycine (equation 12, rate constants from equation 13). Note that *k*std are plotted on a logarithmic scale.

Soil pH is likely a major factor shaping the availability as well as extractability of cyanate because its reactivity is strongly pH-dependent. Cyanate is the anionic form of isocyanic acid, which is a weak acid with a pK_a_ of 3.66, so that cyanate is the dominant species at neutral and alkaline pH (Fig. 2c). Based on what has been observed for other inorganic ions, it is predicted that cyanate adsorption in soils decreases with increasing pH, with no adsorption at pH > 8 (Fig. 2c) (*28*). Such adsorption behavior is in line with the results of our extraction test: at high soil pH, cyanate was completely extracted with water (i.e., no cyanate adsorption), whereas at lower pH (here neutral pH) cyanate extraction was incomplete when extracted with water, but when extracted with salt solutions increasing amounts of cyanate (i.e., exchangeable/adsorbed cyanate) were recovered. In turn, the distinctive low recovery of added cyanate in the acidic soil, as well as the low detectability of cyanate in soils with low pH across a European transect, were most likely due to irreversible reactions of cyanate and in particular isocyanic acid with amino- and carboxylgroups at low pH. Both chemical species hydrolyze abiotically to ammonia/ammonium and on dioxide/bicarbonate in aqueous solution according to three simultaneous reactions, which are strongly pH-dependent: hydronium ion-catalyzed hydrolysis of isocyanic acid (eq. 6; dominant reaction at low pH), direct hydrolysis of isocyanic acid (eq. 7), and direct hydrolysis of cyanate (eq. 8, dominant reaction at high pH). Combining these reactions, the rate of cyanate/isocyanic acid hydrolysis substantially increases with decreasing pH, rendering cyanate unstable at low pH (markedly at pH < 4; Fig. 2d). Moreover, isocyanic acid also reacts with carboxyl, sulfhydryl, phosphate, thiol or phenol groups, which mostly occurs at low pH (*29*).

At neutral to alkaline pH, the most relevant abiotic reactions of cyanate/isocyanic acid in the (soil) environment are the irreversible reaction of isocyanic acid with the amino group of amino acids and proteins (eq. 12; carbamoylation) and the reaction of cyanate and ammonium to urea (eq. 3; equilibrium reaction that favors urea more than 99%). As the rates plotted in Fig. 2d are standardized rates, they do not take into account the concentrations of the two reactants involved in the second order reactions (cyanate and amino acids or cyanate and ammonium). Therefore, the actual rates will depend on the soil solution concentrations of both reactants. Concentrations of amino acids and ammonium in the soil solution are also modulated by their adsorption behavior (i.e., weak or strong), which strongly depends on their chemical properties and on physicochemical properties of the soil, such as clay content and cation exchange capacity (*30*). Therefore, the rates of abiotic reactions of cyanate with amino acids/proteins or with ammonium may strongly vary between different soil types, depending on soil physicochemical properties other than soil pH. For example, low-nutrient soils with high adsorption capacity for ions and low contents of amino acids and ammonium have the greatest potential to limit these abiotic reactions of cyanate. Nevertheless, cyanate is significantly more stable in soils with high pH, as the rate of abiotic hydrolysis of cyanate to ammonium at pH < 4 is about two orders of magnitude higher compared to the reactions with amino acids or ammonium (note that the standardized rates are plotted on a logarithmic scale in Fig. 2d).

### Soil cyanate dynamics

Understanding environmental dynamics and turnover of cyanate requires the knowledge about both pool sizes and fluxes. Therefore, we thoroughly assessed cyanate fluxes in neutral/alkaline soils, where it does not rapidly decompose to ammonium, by using two different approaches: first, we determined the half-life (*t*_1/2_) of cyanate by amending two soils with isotopically labelled cyanate solution (^13^C^15^N-KOCN) and measuring the decrease in concentration over time. To assess abiotic reactions that may limit cyanate bioavailability in neutral/alkaline soils, we also differentiated between biotic and abiotic decomposition processes of cyanate in this approach using sterilized (autoclaved) soils, where enzymatic activities are strongly reduced. Second, we assessed urea quantitively as a source for cyanate formation in soils, by combining an empirical and modelling approach to obtain estimates of gross cyanate production and consumption in a urea-amended soil. Throughout the following discussion, we will refer to these two experiments as “tracer experiment” and “urea addition experiment”, respectively.

In the tracer experiment, we added isotopically labelled cyanate to two distinct soils with the same pH (7.4 in 0.01M CaCl_2_) and similar *in situ* cyanate concentrations: a grassland and an arable soil with soil cyanate concentrations of 27.3 (± 4.7 SE) and 21.2 (± 4.5 SE) pmol g^-1^ soil d.w., respectively. We found that the depletion of isotopically labelled cyanate was substantially faster in the grassland soil than in the arable soil: 58 (± 2 SD) and 25% (± 4 SD) of the labelled cyanate were lost in the grassland and arable soil, respectively, after 90 min of incubation. Here, the depletion of cyanate includes both biotic and abiotic processes. To distinguish abiotic reactions and biotic cyanate consumption over time, we corrected these data for abiotic cyanate loss rates inferred from sterile (autoclaved) soil samples. We then fitted a first order exponential decay curve and used the exponential coefficient to calculate the biotic half-life of cyanate. We found that the grassland soil had a biotic half-life of 1.6 h, which is significantly shorter than that of the arable soil, which was 5.0 h (*t* = 6.64, P < 0.01; Fig. 3). This biotic-mediated turnover of the soil cyanate pool was relatively fast and in the same range as the turnover of free amino acids in soils and plant litter (< 6h) (*31, 32*) and soil glucosamine (*33*). By contrast, mean residence times of soil ammonium and nitrate are found to be around 1 day (half-life of 16.6 h), but can also be in the range of several days due to lower input rates and larger pool sizes. For instance, in arable soils ammonium and nitrate had mean residence times between 0.6 and 7.9 day (half-life of 10.0 h to 5.5 d), and between 1.1 and 25.7 d (half-life of 18.3 h to 17.8 d), respectively (*34*). The abiotic half-life of cyanate determined in sterile soil samples was similar for both soils (*t* = 0.13, P = 0.9024), with 13.4 h and 15.1 h for the grassland and the arable soil, respectively (Fig. 3). The ratio of the biotic (*k*_b_; min^-1^) and abiotic (*k*_a_; min^-1^) rate constant of cyanate consumption was 8 (*k*_b_/*k*_a_ = 0.007/0.0009) for the grassland soil and 3 (*k*_b_/*k*_a_ = 0.002/0.0008) for the arable soil. This shows that the consumption of cyanate in these neutral/alkaline soils is mainly biotic, with only small contributions from abiotic processes.

**Fig. 3.**
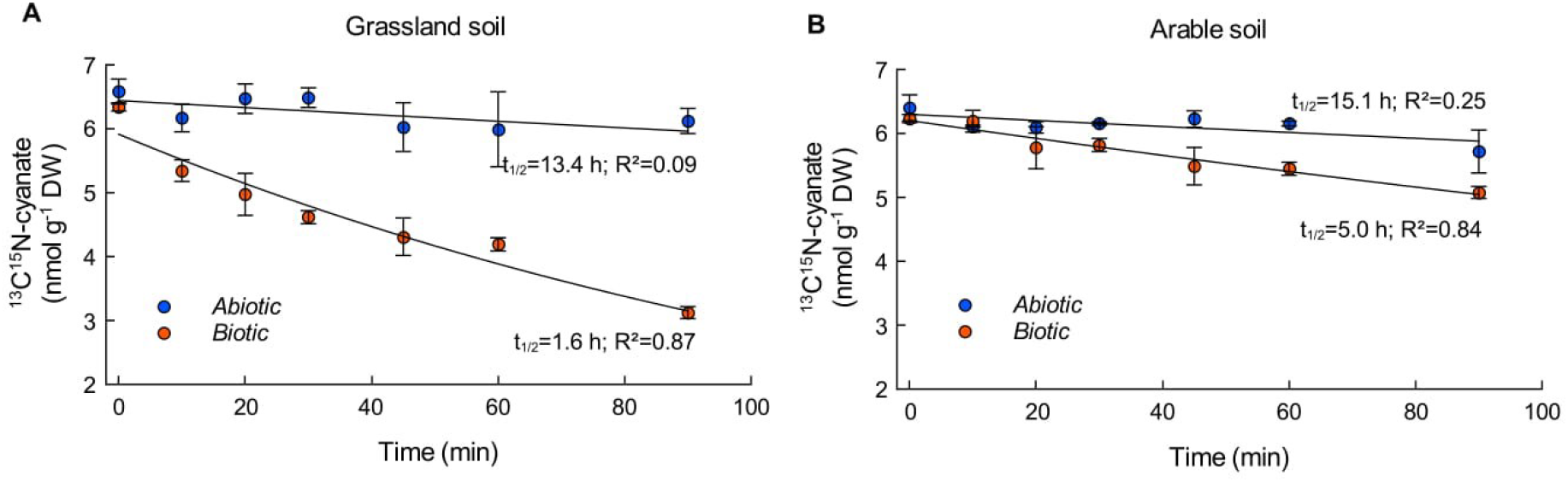
Dynamics of soil ^13^C^15^N-cyanate consumption in two contrasting neutral soils (pH = 7.4), (**A**) a grassland soil and (**B**) an arable soil (“tracer experiment”). ^13^C^15^N-cyanate was added to sterile (i.e., abiotic control) and non-sterile soils, and incubations were stopped after 0, 10, 20, 30, 45, 60 and 90 min. To obtain biotic cyanate consumption over time, the non-sterile samples were corrected for abiotic loss of cyanate derived from the sterile samples. Dynamics of cyanate consumption over time for the corrected non-sterile soils and sterile soils were described by fitting a first order exponential decay curve and the exponential coefficient was used to calculate half-life (*t*_1/2_) of the ^13^C^15^N-cyanate pool. Shown are average values ±1SE (n = 3).

The contribution of urea to soil cyanate formation has never been quantified, although it has been speculated that cyanate formation is the reason for the observed negative effects of urea fertilizer (when applied at high rates) on early plant growth (*35*). It was found that cyanate was toxic to plant cells, although when cyanate was added to soil, it did not have a negative effect on seed germination and plant biomass yield (*35, 36*). Nevertheless, it is unclear whether cyanate accumulates during fertilizer application, and urea-derived cyanate has never been considered in the context of microbial nutrient cycling in agricultural soils. Studying cyanate formation from urea fertilizer application in soils has been hindered by the lack of sensitive analytical methods to measure cyanate in the environment, which has only recently become available (*27*). This is also complicated by the fact that rates of cyanate formation from urea in soils depend on the pool sizes of different N species, which, in contrast to sterile aqueous solutions under laboratory conditions, change over time. These changes are due to microbial activity, i.e., decrease in urea concentration due to ureolytic activity, net change in ammonium concentration as a result of the production from urea hydrolysis and organic matter mineralization, and the consumption and/or immobilization by nitrification, assimilation and soil fixation (abiotic immobilization by clay and humic substances), and the biotic consumption of cyanate.

In order to obtain estimates of gross rates of cyanate dynamics, we developed an approach that combines experimental data and modelling. The chemical equilibrium reaction of urea and ammonium cyanate has been intensively studied and the rate constants for this reaction in aqueous solution are well established under controlled laboratory conditions (eq. 3–5). We took advantage of these well-established rate constants by using them to compute rates of cyanate production and consumption based on observed changes in pool sizes in soil solution (eq. 14 and Fig. 4a). We assume that net changes in cyanate concentration are the result of the production from urea and the biotic and abiotic consumption of cyanate, and that no cyanate adsorption occurs in the alkaline soil used in this experiment.

**Fig. 4.**
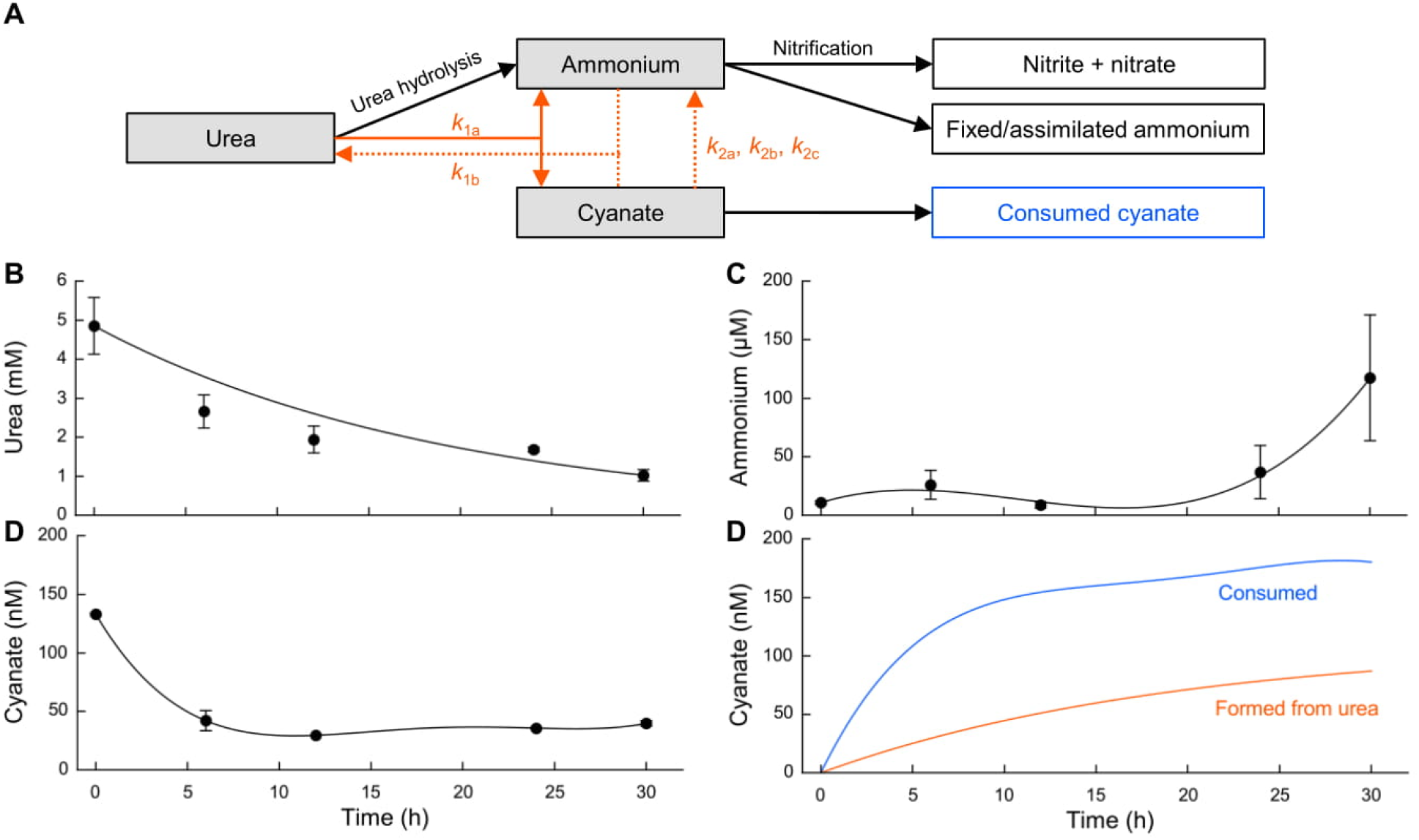
Gross cyanate production and consumption in soil solution of a urea-amended arable soil (“urea addition experiment”). (**A**) Schematic representation of pools and fluxes used to model rates of abiotic cyanate formation from urea and microbial consumption of soil cyanate. Urea, ammonium and cyanate, which are involved in the chemical equilibrium reaction, are highlighted as grey boxes. Rate constants of abiotic reactions are depicted in orange and were used to model cyanate fluxes based on observed pool sizes. We included abiotic hydrolysis of cyanate to ammonium, as the rate constants for the reaction are well established. Panels (**B-D**) show urea, ammonium and cyanate concentrations in soil solution, respectively. Filled circles are observed data (average ± 1SE) at 0, 6, 12, 24 and 30 h after urea addition. (**E**) Modelled rates of gross cyanate production from urea (orange line; eq. 14 using rate constants from eq. 4, 5 and 9–11) are shown as cyanate accumulation over time and gross cyanate consumption (blue line) calculated as the difference between cyanate production and the observed net change in concentration.

For this “urea addition experiment” we used the same arable soil as in the tracer experiment, which was cultivated with rice every second year and received N fertilizer in the form of urea. Urea solution corresponding to the fertilizer application rate of this soil (i.e., 180 kg N ha^-1^ y^-1^) was added, and soil solutions were obtained at several time points throughout a 30-h incubation period. We found that urea was almost completely hydrolyzed at the end of the incubation (Fig. 4b), and that only a very small fraction (<1%) of the resulting ammonium was recovered in soil solution throughout the incubation (Fig. 4c). Thus, most of the ammonium was adsorbed, abiotically fixed, converted to nitrate or assimilated. When urea was added to the soil incubations at the beginning, a small cyanate amount was added along with it. This was unavoidable as cyanate was immediately formed upon urea dissolution when the solution was prepared. This cyanate pool was rapidly consumed during the first 6 h, after which steady cyanate concentrations were reached, indicating balanced production and consumption rates (Fig. 4d). The rate of cyanate formation from urea depends on the pool size of urea, ammonium and cyanate, which change over time (i.e., decrease of urea concentration due to ureolytic activity, while net changes in ammonium concentration are the result of the production from urea hydrolysis and the consumption and ammonium immobilization by nitrification and fixation/assimilation, respectively). For the model, urea concentration over time was described by a first order reaction (eq. 15), and ammonium and cyanate concentrations were fitted with a third and fourth degree polynomial function, respectively (eq. 16 and 17, respectively). By integrating dynamics of biological processes into the abiotic equilibrium reactions of urea (eq. 14), our model estimates cyanate production of 86.8 nM from urea (180 kg N ha^-1^) after 30 h (Fig. 4e), which equals to an average gross cyanate production rate from urea of 2.9 nM h^-1^. Gross cyanate consumption was 6.0 nM h^-1^ (180 nM during 30 h), encompassing also the consumption of the added cyanate through urea addition at the beginning of the incubation. Our study therefore demonstrates that cyanate formed by isomerization of urea was rapidly depleted by soil microorganisms and by abiotic reactions, limiting cyanate accumulation in soils and, thus preventing possible phytotoxic effects of urea-derived cyanate during fertilizer application. The applied empirical modelling approach provides the first estimates of gross cyanate production and consumption rates from urea in a biological/environmental system.

To better grasp the cyanate consumption potential of soil microorganisms, we compared the rate constant of cyanate consumption from the tracer experiment and urea hydrolysis from the urea addition experiment, as both rates followed first order reaction kinetics (Fig. 3b and Fig. 4b, respectively). In the arable soil used for both experiments, we obtained a rate constant of 0.0032 min^-1^ for (biotic) cyanate degradation and 0.0009 min^-1^ for urea hydrolysis, showing that cyanate consumption was approximately 3.7-fold faster than urea hydrolysis. This indicates that soil microorganisms have a remarkably high potential for cyanate consumption, especially by comparison with the well-known rapid hydrolysis of urea in soils due to high ureolytic activity.

However, knowing how much cyanate is continuously produced *in-situ* in (not urea amended) soils is still unsolved. Soil cyanate concentrations were too low for performing an isotope pool dilution assay to determine gross rates of cyanate production and consumption. We therefore explored *in-situ* gross cyanate production rates by an alternative approach. We used concentrations and mean residence times (MRT) of cyanate in soils to calculate gross cyanate production rates assuming steady-state conditions, i.e., productive and consumptive fluxes are balanced, giving a zero net change in cyanate concentration, for an unamended soil (*flux* = *pool*/*MRT*). For the urea addition experiment, we computed MRTs of cyanate for 6 h-time intervals, which ranged between 3.9 to 20.9 h, with lower MRTs at the beginning of the incubation (Table 1). For the tracer experiment, where we added isotopically labelled cyanate, we calculated half-life of cyanate that includes both abiotic and biotic processes for the arable soil (*t*_1/2_ = 3.6h) and converted it to MRT (*MRT* = *t*_1/2_/0.693), which was 5.2 h (Table 1). This MRT is in the same range as the MRTs computed for the first 12 h of the urea addition experiment. Using the MRT of 5.2 h derived from the tracer addition experiment and the *in-situ* cyanate concentration of this soil (21.2 pmol g^-1^ d.w.), we obtained a gross cyanate production rate of 98.8 pmol g^-1^ d.w. d^-1^. This gross cyanate production rate was approximately 4-times higher than the rate at which cyanate is formed through isomerization of urea (26.0 pmol g^-1^ d.w. d^-1^; Table 1). However, additions of substrates can stimulate consumptive processes and, thus, can lead to an overestimation of fluxes in relation to unamended conditions, which consequently results in lower MRTs. Assuming that the MRT derived from the tracer experiment as well as MRTs computed for the first 12 h of the incubation with urea are underestimated due to the substrate addition, we further calculated conservative estimates of gross cyanate production rates, using MRTs of 24 h (which is similar to the MRT for the end of the incubation with urea, when the initial pulse of cyanate was depleted), and 48 h. This yielded gross cyanate production rates of 21.2 and 10.6 pmol g^-1^ d.w. d^-1^, respectively. These rate estimates are still in the same order of magnitude as the average cyanate gross production rate during the 30-h incubation with urea (26.0 pmol g^-1^ d.w. d^-1^; Table 1). These rates are more than 3 orders of magnitude lower than gross rates of N mineralization and nitrification in soils (*37*) and approximately 1-2 orders of magnitude lower than gross production rates of some organic N compounds from microbial cell wall decomposition in soils (*33*). While our calculations do not necessarily represent accurate estimates of *in-situ* gross cyanate production rates, they provide a first approximation of their magnitude in soils, as environmental cyanate production rates are entirely unknown. Most importantly, our data thus suggest that cyanate in unamended soils may be produced at rates similar to rates of cyanate formation from urea fertilizer.

**Table 1.** Estimates of mean residence time (MRT) of cyanate obtained from two approaches, the urea addition and the tracer experiment. We computed MRTs of cyanate and gross cyanate production rates for 6h-time intervals of the urea addition experiment. For comparative analysis of the rates, we converted them from nmol L^-1^ soil solution to rates based on a dry soil mass basis. We used MRTs to calculate gross cyanate production rates for unamended soils, assuming steady-state conditions, i.e., production and consumption fluxes are balanced, resulting in no change in cyanate concentration (*flux* = *pool/MRT*).

|  | MRT<br>(h) | Gross cyanate production<br>(pmol g<sup>-1</sup> dw d<sup>-1</sup>) |
| --- | --- | --- |
| Urea addition experiment (0-30 h) |  | 26.0 |
| Time interval 0-6 h | 3.9 | 39.1 |
| Time interval 6-12 h | 6.5 | 28.9 |
| Time interval 12-18 h | 20.9 | 21.1 |
| Time interval 18-24 h | 19.1 | 15.4 |
| Unamended soil | 5.2* | 98.8‡ |
|  | 24† | 21.2‡ |
|  | 48† | 10.6‡ |
|  | 72† | 7.1‡ |

\*Estimate from tracer addition experiment
†Higher MRTs assumed for conservative calculations
‡Calculated using MRT assuming steady-state conditions of cyanate in soil solution

Sources of cyanate in natural ecosystems are not well understood. It is possible that, in natural/uncontaminated soils, cyanate is formed from cyanide, which can be released by cyanogenic bacteria, fungi and plants into the soil (*38, 39*). Another source of cyanate can be urea excreted by soil fauna or released by lysed microbes. Soil urea concentrations are in the low nmol g^-1^ range (*40*), being about 3 orders of magnitude higher than soil cyanate concentrations. Furthermore, within living organisms, cyanate may result from the non-enzymatic decomposition of carbamoyl phosphate, a nucleotide precursor (*23*), which may leak into the environment during growth or lysis of an organism. It has been shown that net cyanate production occurred in diatom cultures during the stationary phase, but not in a cyanobacterial culture (*22*). However, the pathway of cyanate production in these diatom cultures is unknown. This certainly warrants future work, especially because cyanate production through the repetitive process of organisms’ growth and death would provide a continuous source of cyanate in the environment.

### Cyanate availability across different environments

The cyanate concentrations measured in the soils studied here were low compared to other N pools. The abundance of cyanate was about 3 orders of magnitude lower than ammonium or nitrate in the soils across a European transect. To determine if cyanate concentrations are exceptionally low in soils in general, we compared cyanate concentrations across different environments. As cyanate concentrations are largely unknown in other environments, we analyzed cyanate in salt marsh sediments including pore water, and activated sludge as well as discharge from municipal wastewater treatment plants. We additionally collected published data on marine cyanate concentrations (*22*). As direct comparisons of cyanate concentrations are not possible due to different matrices (seawater, soil extracts, pore water), we normalized cyanate concentrations by calculating ammonium-to-cyanate ratios. Ammonium is a major N source in the environment and can be used as an indicator of the N status of an ecosystem, and, thus, this ratio can be interpreted as a proxy of relative cyanate-N availability. The median of ammonium-to-cyanate ratios was 955 for soil extracts, 1842 for salt marsh sediment extracts, 606 for pore water extracted from salt marsh sediments, 2189 and 514 for activated sludge and discharge of wastewater treatment plants, respectively, and 14 for seawater (Fig. 5). Despite large differences between median values between some environments, we found no significant differences in relative cyanate availability between soils and any of the other environments, except for seawater, which had lower ammonium-to-cyanate ratios (Kruskal-Wallis test followed by Dunn’s test, H(2) = 101.1, P < 0.001). These results indicate that relative cyanate concentrations in soils are similar to those in salt marsh sediments or activated sludge from wastewater treatment plants. Seawater showed the lowest ammonium-to-cyanate ratios, which were significantly lower than for all other environments. Cyanate concentrations in seawater are in the nanomolar range, which is in the same order of magnitude as ammonium concentrations typically found in oligotrophic marine environments (*22, 27, 41*). In contrast to the low MRT of cyanate in soils, that of cyanate in marine surface water has been shown to range between 2.3 d and 8.1 d (similar to MRT of ammonium) but can be as high as 36 d (*41*). Therefore, in marine systems relative concentrations of cyanate are higher but cyanate turnover rates are slower than in terrestrial systems.

**Fig. 5.**
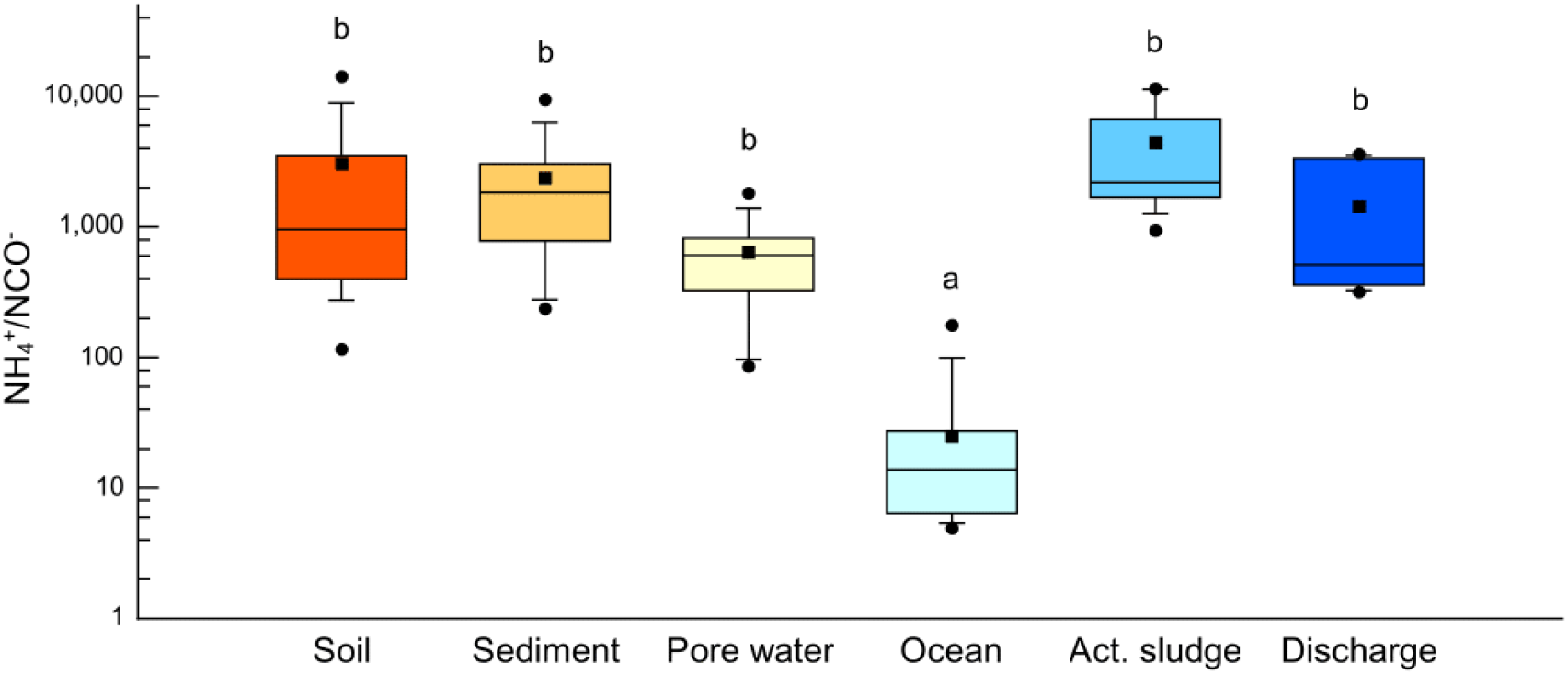
Comparison of relative cyanate availability across different environments. Samples include soils (n=17), salt marsh sediments (n=12), pore water of salt marsh sediments (n=10), ocean (n=75), activated sludge (n=12) and discharge (n=9) from municipal wastewater treatment plants. Relative cyanate availability is represented as the ratio of extractable ammonium over cyanate. Different letters indicate significant differences in relative cyanate availability between environments (Kruskal-Wallis test followed by Dunn’s test). The box plot shows the median (solid line within box), the average (rectangle), 25th and 75th percentiles as vertical bars, 10th and 90th percentiles as error bars and minimum and maximum as circles. Data on marine cyanate and ammonium concentrations are from Widner *et al.* (*22*).

## Conclusion

Soil is a heterogenous environment in regard to its physicochemical properties, and thus assessing cyanate bioavailability requires a thorough analysis of the abiotic and biotic behavior of cyanate. Although neutral/alkaline soil pH favors cyanate stability, it may also be interesting to specifically look at low pH soils with detectable cyanate concentrations, as the faster abiotic decomposition needs to be compensated by higher production rates. Although soil cyanate concentrations may seem quantitatively insignificant compared to those of ammonium, cyanate may constitute an important, yet largely overlooked, N and energy source for soil microorganisms, specifically when considering the relatively high production rates. Additionally, cyanate is more mobile in soil solution compared to ammonium, the availability of which is strongly limited in soils through adsorption, favoring the relative availability of cyanate-N in soil solution. Using cyanate directly as a source of energy, carbon dioxide or nitrogen could thus represent a selective advantage for specific microbial taxa. The ability to use cyanate as a source of reductant (i.e., ammonia) and carbon (i.e., carbon dioxide) may also be an important ecological adaptation of ammonia-oxidizing microorganisms, with implications for soil nitrification. Although only a few genomes of ammonia-oxidizing archaea and complete ammonia-oxidizing (comammox) organisms are known to encode cyanases (*5, 42, 43*), another but yet unknown enzyme may be involved in the decomposition of cyanate. Kitzinger et al. (*7*) found that an isolate of a marine ammonia-oxidizing archaeon lacking a cyanase can oxidize cyanate to nitrite. Furthermore, consortia of cyanase-encoding nitrite-oxidizers and non-cyanase encoding ammonia oxidizers can collectively thrive on cyanate as energy source (*4*). Clearly, the fate of cyanate-N in soils needs to be further investigated, together with the microbial populations that are involved in cyanate turnover or are able to use cyanate directly as a N and energy source. Our study provides a first insight into cyanate dynamics in soils, providing evidence that cyanate is actively turned over in soils and represents a small but continuous N source for soil microbes.

## Materials and Methods

### Cyanate analysis

To test soil extractants for cyanate analysis, three soils (0-15 cm depth) differing in soil pH were collected in Austria, sieved to 2 mm and stored at 4°C. An alkaline grassland soil was collected in the National Park Seewinkel (47° 46’ 32” N, 16° 46’ 20” E; 116 m a.s.l.), a neutral mixed forest soil in Lower Austria (N 48° 20’ 29” N, 16° 12’ 48” E; 171 m a.s.l.) and an acidic grassland soil at the Agricultural Research and Education Centre Raumberg-Gumpenstein (47° 29’ 45” N, 14° 5’ 53” E; 700 m a.s.l.). The recovery of cyanate was assessed by using cyanate-spiked (15 nM potassium cyanate added) and unspiked extraction solutions. We used water (Milli-Q, >18.2 MOhm, Millipore), 10 mM CaSO_4_ and 1 M KCl as extractants. The three soils (n=4) were extracted using a soil:extractant ratio of 1:10 (w:v), shaken for 10 min, and centrifuged (5 min at 14000 × *g*). The supernatant was stored at −80°C until analysis, as it has been shown that cyanate is more stable at −80°C compared to −20°C (*27*).

To explore soil cyanate concentrations across different soil and land management types, and along a climatic gradient, we collected 42 soils from Europe. Sites ranged from Southern France to Northern Scandinavia and included forests (F), pastures (P), and arable fields (A) (Fig. 2a). At each site five soil cores (5 cm diameter, 15 cm depth) were collected, after removal of litter and organic horizons. Soil samples were shipped to Vienna and aliquots of the five mineral soil samples of each site were mixed to one composite sample per site and sieved to 2 mm. In addition to those 42 samples, we collected a rice paddy soil in Southern France (sample code A1; four replicates) and three grassland soils (G) in close vicinity of Vienna, Austria (G1 and G2 from saline grassland, three replicates; G3, one soil sample). Soil samples were stored at 4°C and extracted within a few days. All sampling sites with their location, soil pH, and cyanate, ammonium and nitrate concentrations are listed in Table S1. For cyanate and ammonium analysis, soils (2 g fresh soil) were extracted with 15 mL 1 M KCl, shaken for 30 min and centrifuged (2 min at 10000 × *g*). The supernatants were transferred to disposable 30 mL syringes and filtered through an attached filter holder (Swinnex, Millipore) containing a disc of glass microfiber filter (GF/C, Whatman). To reduce abiotic decay of cyanate to ammonium during extraction, the extraction was performed at 4°C with the extracting solution (1 M KCl) cooled to 4°C prior to extraction. Soil extracts were stored at −80°C until analysis.

To compare cyanate availability across different environments, we analyzed cyanate in salt marsh sediments and activated sludge from municipal wastewater treatment plants, and, additionally, we collected published data on cyanate concentrations in the ocean. We collected sediment samples (0-10 cm, n=4) from a high and low salt marsh dominated by *Spartina alternflora* in New Hampshire, USA (43° 2’ 26” N, 70° 55’ 36” W), and from a *S. alternflora* and a *S. patens* salt marsh in Maine, USA (43° 6’ 31” N, 70° 39’ 56” W). We chose these types of salt marsh because they have been shown to accumulate cyanide (*44*), which potentially could be oxidized to cyanate. Sediment samples were stored at 4°C and extracted within a few days after collection using 2 M KCl at a sediment:extractant ratio of 1:10 (w:v) for 30 min at room temperature. The supernatants were filtered through glass microfibre filters as described above for soil samples. Pore water was extracted with Rhizon samplers (Rhizon CSS, 3 cm long, 2.5 mm diameter, Rhizosphere Research Products, Netherlands) with a filter pore size of 0.15 μm. Triplicate samples of activated sludge were collected from four municipal Austrian wastewater treatment plants (WWTPs), i.e., from Alland (48° 2’ 30” N, 16° 6’ 1” E), Bruck an der Leitha (48° 2’ 4” N, 16° 49’ 7” E), Wolkersdorf (48° 21’ 31” N, 16° 33’ 31” E) and Klosterneuburg (48° 17’ 39” N, 16° 20’ 30” E). Samples from the discharge were also collected from the first three listed WWTPs. Samples were cooled on gel ice packs during the transport to Vienna. Upon arrival in Vienna, samples were transferred to disposable 30 mL syringes and filtered through an attached filter holder (Swinnex, Millipore) containing a disc of glass microfiber filter (GF/C, Whatman). All samples were immediately stored at −80°C until analysis.

Cyanate concentrations were determined using high performance liquid chromatography (HPLC) with fluorescence detection, after conversion to 2,4(1//,3//)-quinazolinedione (*27*). Briefly, a 230 μL aliquot of the sample was transferred to a 1.5 mL amber glass vial, 95 μL of 30 mM 2-aminobenzoic acid (prepared in 50 mM sodium acetate buffer, pH = 4.8) were added, and samples were incubated at 37°C for 30 min. The reaction was stopped by the addition of 325 μL of 12 M HCl. Standards (KOCN) were prepared fresh daily and derivatized with samples in the same matrix. Derivatized samples were frozen at −20°C until analysis. Just before analysis samples were neutralized with 10 M NaOH. The average detection limit was 1.2 nM (± 0.2 SE). Ammonium concentrations were quantified by the Berthelot colorimetric reaction. As direct comparison of cyanate concentrations was not possible across the different environments and matrices, we normalized cyanate concentrations relative to ammonium concentrations, by calculating ammonium-to-cyanate ratios. Data on marine cyanate and ammonium concentrations were taken from Widner et al. (*22*). For marine samples where cyanate was detectable but ammonium was below detection limit, we used the reported limit of detection of 40 nM for ammonium. The presented soil and sediment data are biased towards higher cyanate availabilities (i.e., low NH_4_^+^/NCO^-^ ratios), due to the exclusion of samples where cyanate was possibly present but was below detection limit. Soil pH was measured in 1:5 (w:v) suspensions of fresh soil in 0.01 M CaCl_2_ and water.

### Dynamics of cyanate consumption in soil using stable isotope tracer

For the determination of half-life of cyanate, we used two soils: a grassland soil (G3) and a rice paddy soil (A1). Both soils had a pH of 7.4 (determined in 0.01 M CaCl_2_). The grassland soil had a soil organic C content of 3.7%, soil N content of 0.192%, molar C:N ratio of 22.4, ammonium concentration of 5.60 nmol g^-1^ d.w., nitrate concentration of 1.03 μmol g^-1^ d.w., and an electrical conductivity of 82.0 mS/m. The rice paddy soil had a soil organic C content of 1.0%, soil N content of 0.098%, molar C:N ratio of 11.9, ammonium concentration of 2.47 nmol g^-1^ d.w., nitrate concentration of 0.91 μmol g^-1^ d.w., and an electrical conductivity of 21.7 mS/m. To equilibrate soil samples after storage at 4°C, soil water content was adjusted to 55% water holding capacity (WHC) and soils incubated at 20°C for one week prior to the start of the experiment. To correct for abiotic reactions of cyanate, a duplicate set of soil samples was prepared and one set of them was sterilized by autoclaving prior to label addition while the other set was left under ambient conditions. Soil samples were autoclaved three times at 121°C for 30 min with 48 h-incubations at 20°C between autoclaving cycles to allow spores to germinate prior to the next autoclaving cycle and to inactivate enzymes (*45*).

Preliminary experiments indicated rapid consumption of added cyanate. Thus, to avoid fast depletion of the added cyanate pool, we added 5 nmol ^13^C^15^N-KOCN g^-1^ f.w. (^13^C: 99 atom%; ^15^N: 98 atom%), which equals to approximately 250-fold the *in-situ* cyanate concentration. With the tracer addition the soil water content was adjusted to 70% WHC. After tracer addition, non-sterile and sterile soil samples were incubated at 20°C for a period of 0, 10, 20, 30, 45, 60 and 90 min (n=3) before stopping the incubation by extraction. Soil extractions were performed with 1 M KCl as described above for the 46 soil samples. Soil extracts were stored at −80°C until analysis.

As no method for compound-specific isotope analysis of cyanate existed, we developed a method to measure isotopically labelled and unlabeled forms of cyanate in soil extracts using hydrophilic interaction chromatography coupled to high-resolution electrospray ionization mass spectrometry (HILIC-LC-MS). For this analysis, cyanate was converted to 2,4(1*H*,3*H*)-quinazolinedione as described above for the RP-HPLC method but with some modifications. Aliquots of 280 μL of each sample were transferred to 2 mL plastic reaction vials, and 20 μL of internal standard solution (4 μM ^13^C-KOCN, 98 atom%) were added. To start the reaction, 120 μL of 30 mM 2-aminobenzoic acid (prepared in ultrapure water) were added, and samples were incubated at 37°C for 30 min. The reaction was stopped by the addition of 420 μL 12 M HCl. To remove HCl and bring the target compound into an organic solvent that can be easily evaporated, we performed liquid-liquid extractions using a mixture of ethyl acetate/toluene (85/15 (v/v)). Each sample was extracted 3 times with 1 mL organic solvent mixture. For extraction, samples were thoroughly mixed by vortexing and the tubes were briefly spun down to separate the two phases. The organic phases of each extraction were combined in a 10 mL amber glass vial and dried under a stream of N2. Before analysis, samples were redissolved in 200 μL mobile phase. Samples were analyzed on a UPLC Ultimate 3000 system (Thermo Fisher Scientific, Bremen, Germany) coupled to an Orbitrap Exactive MS (Thermo Fisher Scientific). 2,4(1*H*,3*H*)-quinazolinedione was separated using an Accucore HILIC column (150 mm × 2.1 mm, 2.6 μm particle size) with a preparative guard column (10 mm × 2.1 mm, 3 μm particle size; Thermo Fisher Scientific). We used isocratic elution with 90/5/5 (v/v/v) acetonitrile/methanol/ammonium acetate, with a final concentration of ammonium acetate of 2 mM (pH = 8). The sample injection volume was 7 μL, and the flow rate 0.2 mL min^-1^. The Orbitrap system was used in negative ion mode and in full scan mode at a resolution of 50,000. The source conditions were: spray voltage 4 kV, capillary temperature 275°C, sheath gas 45 units, and AUX gas 18 units. The instrument was calibrated in negative ion mode before sample acquisition using Pierce LTQ ESI Negative Ion Calibration Solution (Thermo Fisher Scientific). To improve the accuracy of absolute quantification, external calibration was paired with an internal calibrant (^13^C-potassium cyanate) to correct for deviations in liquid-liquid extraction efficiency, ionization efficiency and ion suppression. ^13^C-KOCN (98 atom%) and ^13^C^15^N-KOCN (^13^C: 99 atom%; ^15^N: 98 atom%) were purchased from ICON Isotopes. The mass-to-charge (*m/z*) ratio of unlabeled, ^13^C- and ^13^C^15^N-labelled cyanate was 161.0357, 162.0391, and 163.0361, respectively, and the retention time was 2.2 min. The limit of detection was 9.7 nM.

To obtain biotic cyanate consumption rates, the non-sterile samples were corrected for abiotic decomposition of cyanate derived from the sterile (autoclaved) samples. Dynamics of cyanate consumption over time for the corrected non-sterile soils were then described by fitting a first order exponential decay curve:

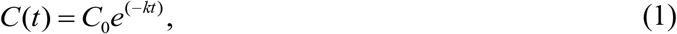

Where *C*(*t*) is the remaining ^13^C^15^N-cyanate concentration at time *t*, *C*_0_ is the initial concentration of ^13^C^15^N-cyanate and *k* is the exponential coefficient for ^13^C^15^N-cyanate consumption. The half-life (*t*_1/2_) of the ^13^C^15^N-cyanate pool was calculated as:

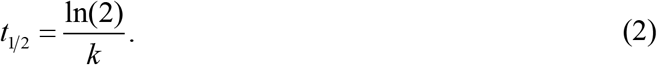

### Abiotic reactions of cyanate and isocyanic acid

Urea (CO(NH_2_)_2_) exists in chemical equilibrium with ammonium cyanate (NH_4_CNO) in aqueous solution:

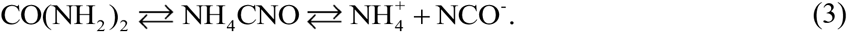

The rate constant for the decomposition of urea (*k*_1a_) and for the conversion of ammonium cyanate into urea (*k*_1b_) were taken from Hagel et al. (*46*), and temperature dependence was calculated by using the Arrhenius equation:

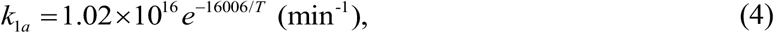

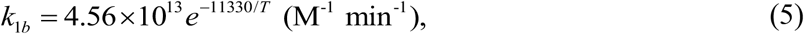

where *T* is temperature in Kelvin.

Cyanate is the anionic form of isocyanic acid. The latter exists as two isomers in aqueous solution, where isocyanic acid is the dominant species. Thus, the acid will be referred to as isocyanic acid. The decomposition of isocyanic acid and cyanate in aqueous solution was found to take place according to three simultaneous reactions:

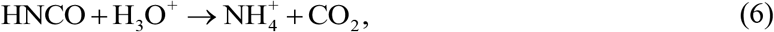

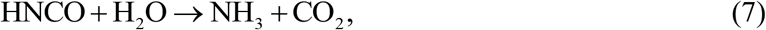

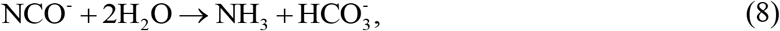

Eq. (6) is for the hydronium ion catalyzed hydrolysis of isocyanic acid (rate constant *k*_2a_; dominant reaction at low pH), eq. (7) is for the direct hydrolysis of isocyanic acid (*k*_2b_), and eq. (8) is for the direct hydrolysis of cyanate (*k*_2c_; dominant reaction at high pH). The rate constants are as follows (*46*):

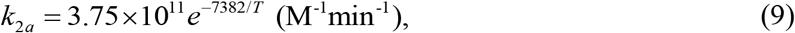

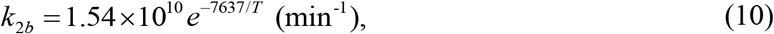

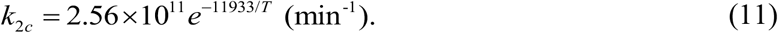

Isocyanic acid reacts with amino groups of proteins, in a process called carbamoylation (*19*):

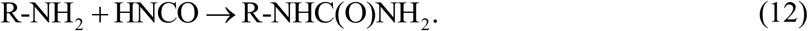

We used glycine as an example for an amino acid, with the following rate constant (*47*):

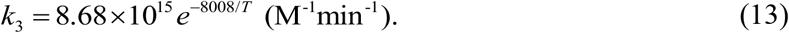

### Urea-derived cyanate formation in a fertilized agricultural soil

For studying the formation and consumption of cyanate after urea addition, we used a rice paddy soil (A1; the same soil as used in the stable isotope tracer experiment), which was cultivated with rice once every second year with a urea application rate of 180 kg N ha^-1^ y^-1^. Treatment of the soil samples was the same as for the stable isotope tracer experiment. Briefly, soil water content was adjusted to 55% water holding capacity (WHC) and soil samples (4 g of fresh soil in a 5 mL centrifugation tube) were incubated at 20°C for one week prior to the start of the experiment. With the addition of the urea solution, the soil water content was adjusted to 70% WHC. We added 140 μg urea g^-1^ soil d.w., which corresponds to approximately 180 kg N ha^-1^. Soil samples were incubated at 20°C for a period of 0, 6, 12, 24 and 30 h (n=4). At each sampling, we collected the soil solution. For this a hole was pierced in the bottom of the 5 mL centrifugation tube containing the soil sample. This tube was then placed into another, intact, 15 mL centrifugation tube and this assembly was then centrifuged at 12000 × *g* for 20 min at 4°C to collect the soil solution. Soil solution samples were stored at −80°C until analysis. For comparative analysis, we converted rates based on nmol/L soil solution to rates based on a dry soil mass basis. For the conversion, we recorded the volume of the soil solution collected and determined the water content of the soil samples after centrifugation.

Cyanate concentrations in soil solution were determined as described above using HPLC. Urea was quantified by the diacetyl monoxime colorimetric method, ammonium by the Berthelot colorimetric reaction and ammonium, and nitrite and nitrate by the Griess colorimetric procedure. For cyanate analysis, aliquots of two replicates were pooled because of insufficient sample volume.

We used the well-established rate constants for the equilibrium reaction of urea in aqueous solution and decomposition of cyanate to ammonia/ammonium and carbon dioxide/bicarbonate, to model gross cyanate production and consumption after urea amendment from observed changes in urea, ammonium and cyanate concentrations over time. Cyanate accumulation was calculated as cyanate formation from urea (rate constant *k*_1a_, eq. 4) minus the conversion of ammonium cyanate into urea (rate constant *k*_1b_, eq. 5), and minus abiotic cyanate hydrolysis to ammonium and carbon dioxide (rate constants *k*_2a_, *k*_2b_, *k*_2c_, eq. 9–11). It has been found that only the ionic species (i.e., NCO^-^ and NH_4_^+^) are involved in the reaction of ammonium cyanate to urea. The difference between cyanate accumulation and the net change in cyanate concentration over time gives then cyanate consumption, as follows:

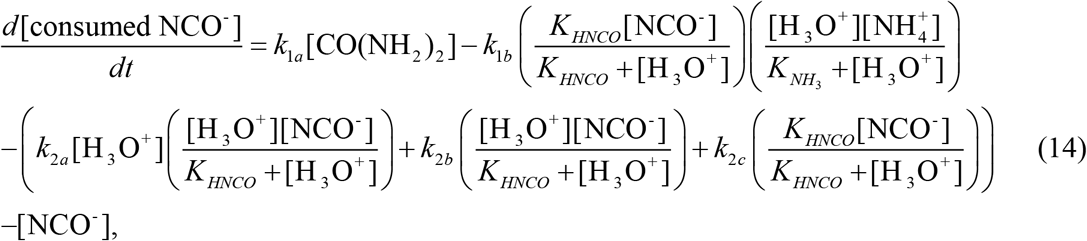

where [NCO^-^] represents the concentration of cyanate and isocyanic acid, [NH_4_^+^] is the sum of ammonium and ammonia, *K*_HNCO_ and *K*_NH3_ is the acid dissociation constant of isocyanic acid and ammonia, respectively, and [H_3_O^+^] is the hydronium ion concentration. Urea concentration over time was described by a first order reaction (eq. 15; unit of rate constant is min^-1^), and ammonium and cyanate concentrations were fitted with a third and fourth degree polynomial function, respectively (eq. 16 and 17, respectively), as follows:

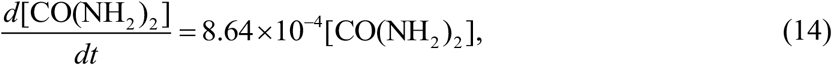

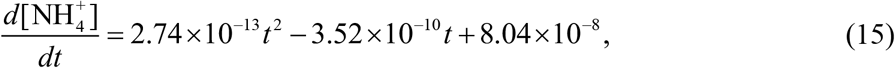

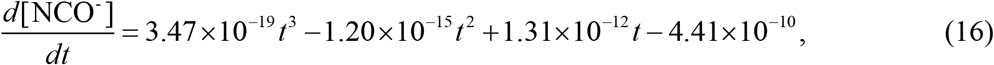

where *t* is time in min and concentrations are mol/L soil solution.

The input parameters were 7.4 for pH (pH of solution: 7.4 ± 0.1 SD) and 20°C for temperature. As rate constant *k*_1b_ is dependent on the ionic strength, we corrected the rate constant (given at *I* = 0.25 (*46*)) using the Extended Debye-Hückel expression:

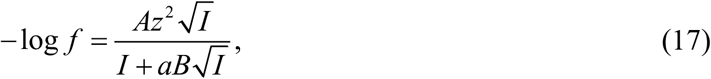

Where *f* is the activity coefficient, *A* and *B* are constants that vary with temperature (at 20°C, A = 0.5044 and B =3.28 × 10^8^), *z* is the integer charge of the ion, and *a* is the effective diameter of the ion (a = 5 Å;, *46*). We used an ionic strength *I* = 0.01, which is within the range observed for soils.

### Statistical Analysis

Statistical significance of the difference between extractants within each soil type was analyzed by one-way analysis of variance (ANOVA) followed by Tukey HSD post-hoc test. For each extractant, statistical significance of the difference between added and recovered cyanate was tested using *t*-test on raw data. To analyze the effect of type of environment on relative cyanate availability (i.e., NH_4_^+^/NCO^-^), we used the Kruskal-Wallis test (assumption for parametric procedure were not met) followed by a non-parametric multiple comparison test (Dunn’s test). For solving differential equations in the model, we used the “deSolve” package in R (*48*).

## Supplementary Materials

Table S1. All soil sampling sites with their location, soil pH, and cyanate, ammonium and nitrate concentrations.

## Acknowledgments

We thank Ricardo J. E. Alves for helpful comments on the manuscript. We thank Ludwig Seidl for assistance with HPLC, Yuntao Hu for help with LC-MS, Roland Albert and Margarete Watzka for collecting soil samples at the National Park Seewinkel, Cyrille Thomas from the Centre Français du Riz, France, for providing soil samples, Markus Schmid for help with collecting samples from wastewater treatment plants, and Lisa Noll, Qing Zheng, Shasha Zhang, Yuntao Hu and Daniel Wasner for collecting soil samples across Europe and providing data on soil pH. We are grateful to the National Park Seewinkel, Austria, and to the wastewater treatment plants in Alland, Bruck an der Leitha, Klosterneuburg and Wolkersdorf, Austria, for permission to collect samples.

## Funding

This study was supported by European Research Council Advanced Grant project NITRICARE (294343).

## Author contributions

M.M. and M.W. designed the experimental concept; M.M. performed experimental work, data analysis and modelling; M.M. developed analytical tools with advice of W.W.; S.J., W.W., and A.R. provided resources and samples. All authors contributed to data interpretation. The manuscript was written by M.M. with input from all authors.

## Competing interests

The authors declare that they have no competing interests.

## Data and materials availability

All data needed to evaluate the conclusions in the paper are present in the paper and/or the Supplementary Materials. Additional data related to this paper may be requested from the authors.

## Notes

### Competing Interest Statement

The authors have declared no competing interest.

