## Supplementary Table 1 for "Cyanate – a low abundant but actively cycled nitrogen compound in soil"

### Supplementary Materials

**Table S1. All soil sampling sites with their location, soil pH, and cyanate, ammonium and nitrate concentrations.** n.d., not determined; b.d., below detection;

| ID | Land use | Latitude | Longitude | pH $H_2O$ | pH $CaCl_2$ | Cyanate pmol $g^{-1}$ | $NH_4^+$ nmol $g^{-1}$ | $NH_4^+/NCO^-$ | $NO_3^-$ nmol $g^{-1}$ |
|---|---|---|---|---|---|---|---|---|---|
| A1 | Arable | 43.66423 | 4.59687 | 8.4 | 7.4 | 21.2 | 2.5 | 117 | 913.8 |
| F1 | Forest | 47.40590 | 14.19570 | 4.7 | 3.5 | b.d. | 104.3 | | 56.9 |
| P1 | Pasture | 47.40710 | 14.19843 | 5.0 | 4.2 | 19.2 | 135.8 | 7079 | 1148.2 |
| P2 | Pasture | 47.41268 | 14.17830 | 5.7 | 4.9 | 18.2 | 257.9 | 14187 | 1188.5 |
| F2 | Forest | 47.41423 | 14.17893 | 4.4 | 3.3 | b.d. | 170.1 | | 124.8 |
| F3 | Forest | 47.48260 | 14.09368 | 4.4 | 3.6 | b.d. | 30.6 | | 25.5 |
| A2 | Arable | 47.49623 | 14.10073 | 6.4 | 5.4 | b.d. | 15.6 | | 697.3 |
| P3 | Pasture | 47.49628 | 14.10072 | 5.5 | 4.6 | 34.9 | 122.2 | 3498 | 812.8 |
| G1 | Grassland | 47.77556 | 16.77222 | n.d. | 7.9 | 93.7 | 63.8 | | 159.4 |
| G2 | Grassland | 47.77556 | 16.77222 | n.d. | 8.0 | 131.1 | 88.2 | | 260.6 |
| G3 | Grassland | 48.35072 | 16.26183 | 8.6 | 7.4 | 27.3 | 5.6 | 205 | 1027.8 |
| P4 | Pasture | 48.37188 | 9.45731 | 8.1 | 7.2 | 65.6 | 25.7 | 392 | 2802.6 |
| A3 | Arable | 48.37283 | 9.45771 | 7.9 | 7.1 | 37.2 | 14.9 | 400 | 1819.6 |
| F4 | Forest | 48.38878 | 9.49017 | 5.8 | 4.7 | b.d. | 852.9 | | 439.2 |
| P5 | Pasture | 50.59696 | 12.03651 | 4.9 | 4.0 | b.d. | 131.3 | | 461.3 |
| F5 | Forest | 50.61323 | 12.04417 | 4.0 | 3.0 | b.d. | 521.3 | | 52.1 |
| A4 | Arable | 50.61350 | 12.04198 | 5.6 | 4.9 | b.d. | 18.3 | | 2317.6 |
| P6 | Pasture | 51.12193 | 10.41270 | 7.6 | 6.8 | 29.5 | 28.2 | 955 | 1529.6 |
| F6 | Forest | 51.13165 | 10.36401 | 5.7 | 4.9 | b.d. | 776.1 | | 1049.8 |
| A5 | Arable | 51.14135 | 10.41074 | 5.1 | 4.6 | b.d. | 40.9 | | 6111.1 |
| F7 | Forest | 53.01260 | 13.91104 | 4.3 | 3.5 | b.d. | 136.5 | | 463.7 |
| A6 | Arable | 53.01690 | 13.71915 | 6.5 | 5.7 | 17.5 | 15.3 | 872 | 1440.3 |
| P7 | Pasture | 53.02525 | 13.82253 | 5.7 | 4.8 | b.d. | 136.3 | | 592.3 |
| P8 | Pasture | 54.11688 | 11.83788 | 6.8 | 5.9 | 18.7 | 22.0 | 1176 | 1338.5 |
| A7 | Arable | 54.11692 | 11.83729 | 7.1 | 6.2 | 19.7 | 6.3 | 320 | 904.0 |
| F8 | Forest | 54.13322 | 11.86038 | 4.3 | 3.4 | b.d. | 24.3 | | 625.3 |
| P9 | Pasture | 58.09573 | 15.78310 | 5.7 | 4.6 | b.d. | 18.1 | | 679.3 |
| A8 | Arable | 58.09629 | 15.78172 | 5.9 | 4.8 | b.d. | 19.9 | | 990.2 |
| F9 | Forest | 58.10546 | 15.77125 | 5.0 | 3.9 | 7.8 | 48.0 | 6180 | 21.8 |
| A9 | Arable | 59.05029 | 11.01064 | 7.4 | 6.6 | 9.0 | 12.6 | 1406 | 1126.2 |
| P10 | Pasture | 59.07636 | 10.95093 | 5.4 | 4.3 | b.d. | 24.5 | | 406.3 |
| F10 | Forest | 60.40044 | 15.18288 | 5.1 | 4.0 | b.d. | 200.0 | | 30.8 |
| P11 | Pasture | 60.41601 | 15.36828 | 6.6 | 5.5 | b.d. | 7.5 | | 634.7 |
| A10 | Arable | 60.42217 | 15.33420 | 6.6 | 5.6 | b.d. | 8.7 | | 469.7 |
| F11 | Forest | 63.96400 | 18.06829 | 4.8 | 3.9 | b.d. | 5.7 | | 21.8 |
| P12 | Pasture | 64.07430 | 18.40021 | 5.5 | 4.4 | 9.7 | 113.9 | 11737 | 254.6 |
| F12 | Forest | 65.96376 | 19.69301 | 5.8 | 4.7 | b.d. | 4321.3 | | 119.4 |
| F13 | Forest | 67.15055 | 26.93055 | 4.9 | 3.7 | b.d. | 226.8 | | 68.6 |
| F14 | Forest | 67.92111 | 24.38552 | 4.8 | 3.9 | b.d. | 10.1 | | 26.4 |
| F15 | Forest | 68.16030 | 15.60596 | 4.5 | 3.3 | b.d. | 1844.4 | | 58.2 |
| P13 | Pasture | 68.16876 | 15.55815 | 4.9 | 3.8 | b.d. | 894.0 | | 713.5 |
| F16 | Forest | 68.26011 | 19.28501 | 4.9 | 3.9 | b.d. | 15.7 | | 301.5 |
| F17 | Forest | 68.27478 | 19.26456 | 4.8 | 3.5 | b.d. | 13.1 | | 25.9 |
| F18 | Forest | 68.28161 | 19.26480 | 4.8 | 3.7 | b.d. | 38.3 | | 9.1 |
| F19 | Forest | 68.34471 | 18.83247 | 4.5 | 3.3 | 10.4 | 20.0 | 1935 | 13.9 |
| F20 | Forest | 71.03227 | 25.79955 | 5.7 | 4.4 | b.d. | 247.7 | | 139.5 |
